# Conserved YKL-40 changes in mice and humans after postoperative delirium

**DOI:** 10.1101/2022.09.26.509551

**Authors:** Jennifer David-Bercholz, Leah Acker, Ana I Caceres, Pau Yen Wu, Saanvi Goenka, Nathan O Franklin, Ramona M Rodriguiz, William C Wetsel, Michael Devinney, Mary Cooter Wright, Henrik Zetterberg, Ting Yang, Miles Berger, Niccolò Terrando

## Abstract

Delirium is a common postoperative neurologic complication among older adults. Despite its prevalence of 14-50% and likely association with inflammation, the exact mechanisms underlying postoperative delirium are unclear. This project aimed at characterizing systemic and central nervous system (CNS) inflammatory changes following surgery in both mice and humans. Matched plasma and cerebrospinal fluid (CSF) samples from the “Investigating Neuroinflammation Underlying Postoperative Brain Connectivity Changes, Postoperative Cognitive Dysfunction, Delirium in Older Adults” (INTUIT; NCT03273335) were used to parallel murine endpoints. Delirium-like behavior was evaluated in aged mice using the 5-Choice Serial Reaction Time Test (5-CSRTT). Using a well-established orthopedic surgical model in the FosTRAP reporter mouse, we detected neuronal changes in the prefrontal cortex, an area implicated in attention, but notably not in the hippocampus. In aged mice, plasma interleukin-6 (IL-6), chitinase-3-like protein 1 (YKL-40), and neurofilament light chain (NfL) levels increased after orthopedic surgery, but hippocampal YKL-40 expression was decreased. Given the growing role of YKL-40 in delirium and other neurodegenerative conditions, we assayed human plasma and CSF samples. Plasma YKL-40 levels were also similarly increased after surgery, with a trend toward greater post-operative plasma YKL-40 increase in patients with delirium. In contrast to plasma, YKL-40 levels in CSF decreased following surgery, which paralleled the findings in the mouse brain. Finally, we confirmed changes in blood-brain barrier (BBB) after surgery as early as 9 hours in mice, which warrants for more detailed and acute evaluations of BBB integrity following surgery in humans. All together, these results provide a nuanced understanding of the neuroimmune interactions underlying post-operative delirium in mice and humans, and highlight translational biomarkers to test potential cellular targets and mechanisms.

## 1. Introduction

Delirium is a neurological complication characterized by changes in cognition, attention, and arousal [1]. Aging is the most prominent risk factor for postoperative delirium; however, we currently do not understand the mechanism whereby aging contributes to the onset of delirium. Delirium is observed frequently after routine surgical procedures, such as after orthopedic surgery, which comprises 4 of the 5 most common surgical procedures in US hospitals [2]. After surgery, delirium can affect up to 50% of older adults [3] and the condition adds $33.3 billion to annual US healthcare costs [4], rendering its impact highly significant to patients, families, and overall healthcare.

Delirium is clinically evaluated by neuropsychological testing, with the Confusion Assessment Method (CAM) being the most widely-used research tool [5]. In clinical practice a shorter form of the CAM, the three-minute CAM (3D-CAM), is often used [6]. Like the long-form CAM, the 3D-CAM has been validated against “gold standard” psychiatric interviews [6]. Further, the 3D-CAM has good overall agreement with the long form CAM as it assesses delirium along the same four domains: (1) acute change and/or a fluctuating disease course; (2) inattention; (3) disorganized thinking, and/or (4) altered level of consciousness [6, 7].

Aside from cognitive testing, more recent efforts have been devoted to biomarker discovery with the intent of establishing predictive molecular signatures that can be related to worse postoperative neurocognitive outcomes. Part of this effort is in parallel with the development and validation of animal models to reliably address cellular and molecular mechanisms underlying delirium and other acute perioperative neurocognitive disorders [8].

We have been interested in modeling key features of delirium in rodents and have focused on orthopedic trauma, arthroplasty surgery, various common mechanisms of injury (e.g., falls), and different types of surgery performed in older adults [9]. Using these models, we and others have described a key role for pro-inflammatory cytokines in triggering neuroinflammation, including changes in glial activation (microglia and astrocytes) and cytokines associated with the behavioral features found in patients with delirium. In particular, we have focused on inattention as one of the key clinical domains [10, 11]. In addition, our murine model has revealed changes in blood-brain barrier (BBB) permeability after surgery, a complication that has now been observed in the clinic [12].

YKL-40 (chitinase 3-like 1) is a glycoprotein secreted by multiple cell types including macrophages, vascular smooth muscle cells, astrocytes and microglia, which has been implicated in a variety of cancers and more recently in neurodegenerative disorders [13]. YKL-40 is often upregulated in the CSF of patients with prodromal Alzheimer’s disease (AD) [14], and mild cognitive impairments (MCI), suggesting its putative role as an early “immune” biomarker. However, its upregulation has not been described in other dementias, possibly due to its diffuse role across multiple immune cell-types, including reactive astrocytes. Notably, YKL-40 levels, which are increased in plasma after lower extremity orthopedic surgery in older adults [15], have been proposed as a strong biomarker predictor in plasma for postoperative delirium [16]. However, the origin of plasma YKL-40 after surgery remains unclear. Further, it is unknown whether elevated plasma YKL-40 levels after surgery reflect an increase in systemic and/or central levels of YKL-40.

The present studies are focused on evaluating systemic and CNS changes in YKL-40 in aged mice following surgery. Using a parallel human cohort of 22 non-neurological, non-cardiac surgery patients aged ≥ 60 years from the INTUIT study [17] with matched plasma and CSF samples, we investigated whether responses for YKL-40 were conserved both in mice and humans.

## 2. Methods

### 2.1 Animals

#### 2.1.1. Animal characteristics and care

Adult (17-24 months old) male C57BL/6 mice (The Jackson Laboratory, Bar Harbor, ME) were used in these experiments. In addition. 3-4 month-old male and female Fos-TRAP2;Ai14 mice (gift of Dr. Fan Wang) [18] were also used. All mice were housed 3-5 per cage under controlled temperature and humidity conditions with a 14:10 h light:dark cycle. Mice were fed rodent chow (Prolab RMH3500, Autoclavable; LabDiet, St. Louis, MO) and *ad libitum* access to food and water. All experiments were conducted under an approved protocol from the Institutional Animal Care and Use Committee at Duke University Medical Center and according to the guidelines described in the National Science Foundation “Guide for the Care and Use of Laboratory Animals” (2011).

#### 2.1.2 Fos-TRAP2; Ai14 and 4-OHT administration

The Targeted Recombination in Active Populations (TRAP) system is a tool that creates permanent genetic access to neurons activated by any stimulation [19]. With the recently developed FosTRAP mouse, Cre recombinase is stimulated when neurons are activated in the presence of tamoxifen to produce permanent expression of td-tomato [18]. FosTRAP mice were housed individually with at least two environmental enrichments for approximately 10 days before 4-hydroxytamoxifen (4-OHT) administration day; (Cat# H6278, Sigma-Aldrich, St. Louis, MO, USA). The 4-OHT was dissolved into 100 % ethanol to make a 20 mg/ml stock solution, samples were aliquoted, and frozen at -20°C. On the administration day, an aliquot of the 4-OHT solution was added to sunflower seed oil (Cat #S5007; Sigma-Aldrich, St. Louis, MO, USA), heated to 50°C to remove the ethanol, yielding a final 4-OHT solution of 10 mg/ml. 50 mg/kg 4-OHT (i.p.) was administered 30 minutes prior to mice undergoing orthopedic surgery or to naïve controls. All mice underwent sevoflurane anesthesia (Covetrus, North America, Dublin, OH, USA) during surgery or for non-operated naive controls and were returned to their individual cages, minimizing any source of distress prior to termination.

#### 2.1.3 Murine orthopedic surgery

Tibial facture was performed as described [10]. Mice were anesthetized with isoflurane via a low-flow digital anesthesia system (SomnoSuite apparatus; Kent Scientific Corporation, Torrington, CT, USA). Body temperature was maintained at 36.5 ± 0.6°C using a homoeothermic pad system (Kent Scientific Corporation). The muscles were dissociated following an incision on the left hind-paw. A 0.38-mm stainless steel pin was inserted into the tibial intramedullary canal, followed by osteotomy, and the incision was closed with 6-0 Prolene suture. Naïve controls did not undergo surgery.

#### 2.1.4 Tissue Collection and Immunohistochemistry

##### 2.1.4.1 Dissection and tissue preparation

For immunohistochemistry (IHC), mice were euthanized under deep isoflurane anesthesia via trans-cardiac perfusion with 50 ml ice-cold 0.1 M phosphate-buffered saline (PBS; pH 7.4), followed by 50 ml ice-cold 4% paraformaldehyde in PBS (PFA). Brains were dissected from the skull, placed in 4% PFA overnight, washed the next day with PBS, placed in a 30% sucrose solution for 3 days, and then stored in optimal cutting temperature (OCT) compound at −80LJ°C until sectioning. The post-fixed brains were coronally cut with a cryostat (Leica CM 1950; Leica Biosystems, Deer Park, IL, USA) in 45 μm (IHC) or 80 μm sections (FosTRAP brains).

##### 2.1.4.2 Immunostaining

Free-floating sections were washed 3 times with PBS (15 min each), blocked for 1 hour in blocking buffer (2.5% BSA, 0.3% Triton X-100 PBS), and incubated overnight at 4°C with the primary antibody in the blocking buffer. The following antisera were used: rabbit anti-AQ-4 (1:500, #AB3594; Millipore), rabbit anti-YKL-40 (1:300, #PA5-43746; ThermoFisher Scientific), goat anti-CD31 (1:500, #AF3628; R&D Systems), rabbit anti-NeuN (1:1000, #ABN78; Millipore, -Sigma-Aldrich), and mouse anti-glial fibrillary acidic protein (GFAP; 1:500, #G3893; Sigma-Aldrich). The next day sections were washed 3 times with the blocking buffer (15 min each) and subsequently incubated for 1 hour at room temperature with the secondary antibody in the blocking buffer. The secondary antibodies consisted of donkey anti-rabbit 647 (1:1000, #AB150075; ABCAM), donkey anti-mouse 594, anti-goat 488, and anti-rabbit 488 (each 1:1000, #A21203, #A11055, #A21206, respectively; Invitrogen, and ThermoFischer Scientific). Lastly, sections were washed 2 times with PBS (10 min each) and finally mounted onto slides covered with DAPI mounting media (#F6057; Sigma-Aldrich).

##### 2.1.4.3 Imaging

Image stacks of the regions of interest (ROIs) were acquired with a 20x or 40x objective using the Zeiss inverted 880 confocal microscope. For Fos-TRAP2 samples, the sections were acquired with a Dragonfly 505 spinning-disk confocal microscope with a 10x or 20x objective for imaging. Maximum intensity projections were analyzed using FIJI software by threshold analysis after background subtraction. Automatic algorithms were determined to match the best the cell morphology and to decrease the noise/signal ratio as described previously [20].

### 2.1.5 Tissue Collection, RNA Extraction, cDNA Synthesis, and PCR

#### 2.1.5.1 Tissue collection

Animals were euthanized under deep isoflurane anesthesia via trans-cardiac perfusion with 50 ml ice-cold 0.1 M phosphate buffered saline only. After perfusion, the brains were removed from the skull, and hippocampi were microdissected on a metallic plate on ice, snap-frozen in liquid nitrogen, and stored at −80□°C until further use.

#### 2.1.5.2 RNA extraction and cDNA synthesis

Total RNA was isolated from the tissue homogenates of each sample using the Qiagen RNeasy Lipid Tissue Mini Kit (Qiagen, Germantown, MD, USA). The homogenization of all the tissues was performed in a NextAdvance bullet blender at 4ºC in Eppendorf Safe-Lock tubes. cDNA synthesis was performed with 200 ng RNA immediately after RNA isolation using a High-Capacity Reverse-Transcription Kit (Applied Biosystems, Foster City, CA) following the manufacturer’s instructions. The RNA quality and quantity were determined with a NanoDrop 1000 Spectrophotometer (Qiagen, Germantown, MD, USA) with RIN values above 8.0.

#### 2.1.5.3 Real-time PCR (qPCR)

TaqMan® Gene Expression Assays (Applied Biosystems, Foster City, CA, USA) were used for expression assessment using reactions containing 10 μl of TaqMan® Fast Advanced Master Mix (2x), 1 μl of the specific TaqMan® assay, 5 μl cDNA, and DNAase/RNAase free-water to a 20 μl volume. Thermal cycling was performed on a Real-Time PCR system Light Cycler 480 (Roche, Indianapolis, IN, USA) in 384-well plates. Each reaction was performed in triplicate and normalized to endogenous 18S gene expression. The CT value of each well was determined using the LightCycler 480 software and the mean of the triplicates was calculated. The relative quantification was determined by the ΔΔCT method [21]. TaqMan® Gene Expression Assays used the following: Aqp4: Mm00802131_m1; Ykl40: Mm00801477_m1; 18S rRNA Mm03928990_g1; IL-1β : Mm00434228_m1; Cd31 : Mm01242576_m1 ; ZO1: Mm01320638_m1; Ocln : Mm00500912_m1; Cldn5 : Mm00727012_s1.

#### 2.1.6 5-Choice Serial Reaction Time Test (5-CSRTT)

The 5-CSRTT was conducted as previously described [10], with minor modifications. The food magazine dispensed single food rewards (20 mg sucrose cocoa-based pellet; BioServ, Flemington, NJ, USA) and testing occurred daily with 25 trials/day at a 1.5 sec stimulus duration. Mice were trained in the task, their baseline responses were recorded, and post-surgical testing began one day after surgery or following the naïve control treatment.

### 2.1.7 Plasma Collection and Analyses

#### 2.1.7.1 Plasma collection

Blood was collected from mice *via* cardiac puncture with a heparinized syringe under terminal anesthesia. The blood was centrifuged at 6,500 RPM (Sorvall legend, micro 21R centrifuge, Thermo-Scientific, Waltham, MA, USA) for 10 min at 4°C, and the plasma stored at −80°C before analysis.

#### 2.1.7.2 ELISA

ELISA kits were used to quantitate the mouse plasma levels of YKL-40 (1:40 dilution, #AB238262; ABCAM, Cambridge, MA, USA), IL-6 (1:4-1:2 dilution, #KMC0061; ThermoFisher Scientific, Waltham, MA, USA), and NfL (1:4 dilution, #103186; Quanterix, Middlesex Turnpike, MA, USA). These samples were collected and evaluated at 24 hours after tibial fracture *per* the manufacturer’s instructions.

### 2.2 Human Patients

#### 2.2.1. Subject inclusion and treatment

The 22 human patients (Table 1) were derived from a subset of patients from a larger cohort study: Investigating Neuroinflammation Underlying Postoperative Brain Connectivity Changes, Postoperative Cognitive Dysfunction, Delirium in Older Adults (INTUIT). Briefly, INTUIT is a prospective observational single-site cohort study of patients age ≥60 years of age undergoing non-neurological, non-cardiac surgery for more than a 2 hours scheduled duration. Exclusion criteria included incarceration, inadequate English fluency, and anticoagulant use that would preclude lumbar punctures. INTUIT had no cognitive exclusion criteria. The INTUIT sub-cohort in this analysis was selected according to age and surgery-type match between subjects who were *APOE4* carriers and non-carriers. Hence, the cohort under the present experiment represented an *APOE4* carrier-enriched group relative to the overall INTUIT cohort. The present experiment was approved by the Duke Health Institutional Review Board and was registered on clinicaltrials.gov (NCT03273335) [17]. All participants, or legally-authorized representatives, provided written informed consent prior to study participation.

**Table 1.** Cohort characteristics

|  | **All (n=22)** |
| --- | --- |
| **Age (years), mean (SD)** | 68 (5) |
| **Male sex, n (%)** | 11 (50) |
| **Years of education, median (Q1, Q3)** | 16 (14, 17) |
| **Baseline Mini-Mental Status Exam Score, median (Q1, Q3)** | 28 (25, 29) |
| **APOE4 Carrier, n (%)** | 9 (41) |
| **ASA physical status, n (%)** |  |
| **2** | 7 (32) |
| **3** | 13 (59) |
| **4** | 2 (9) |
| **Surgical service, n (%)** |  |
| **Thoracic** | 4 (18) |
| **General surgery** | 7 (32) |
| **Gynecology** | 3 (14) |
| **Orthopedics** | 4 (18) |
| **Urology** | 4 (18) |
| **Surgery duration (min incision to end procedure), median (Q1, Q3)** | 137 (74, 168) |

#### 2.2.2 Baseline Cognitive Assessment and Delirium Assessment

Subjects underwent a cognitive battery preoperatively, as described [17]. This battery included the mini-mental status exam (MMSE) - a commonly used cognitive screening tool. The 3D-CAM was administered preoperatively to establish a baseline and then was administered twice daily throughout the postoperative hospitalization period to capture the fluctuations in mental status that characterize delirium. The 3D-CAM assessment was scored for both for the presence or absence of delirium, as well as delirium severity using the “raw” scoring method [7].

#### 2.2.3 Human Plasma and CSF collection

Blood samples were collected before and 24 hours after surgery using standard procedures, including a clean puncture or draw-back from an arterial or venous line with waste drawn prior to sample collection to account for fluid in the tubing and to prevent sample dilution. Blood was collected in EDTA-containing tubes and centrifuged at 3,500 RPM for 15 min at 4°C. Plasma was aliquoted into 1.5 ml microcentrifuge tubes (VWR International) using low-binding pipette tips (Genesee Scientific) and stored at −80°C before analysis. CSF samples were collected via lumbar puncture using sterile technique before and 24 hours after surgery, as described [22].

### 2.2.4 ELISA

#### 2.2.4.1 Assay of YKL-40 in plasma

YKL-40 concentrations in plasma (YKL-40, 1:160, ab255719, ABCAM, Cambridge, MA) were quantitated in samples collected prior to surgery and 24 hours post-operative from the same subjects using the manufacturer’s instructions.

#### 2.2.4.2 Assay of YKL-40 in CSF

YKL-40 concentrations in the CSF were quantitated in one round of analyses with one batch of reagents according to the manufacturer’s instructions using a commercially ELISA kit (R&D Systems, Minneapolis, MN) according to instructions from the manufacturer instructions. The assay was conducted by a board-certified laboratory technician who was blinded to the clinical data.

### 2.3 Statistics

Mice data are presented as means and standard errors of the mean (SEMs). Attention task performances collected from the same mice over time were analyzed with repeated-measures ANOVA (RMANOVA) followed by Bonferroni corrected pair-wise comparisons by day. Differences between control and surgical mice were compared via student t-tests. Human data are presented as means and standard deviation of the mean (SD) or median and Quartile 1,3. Biomarker data were analyzed using t-tests for normally distributed data or Wilcoxon rank-sum tests (WRS) for data that failed the Shapiro-Wilk test for normality. The mouse data were analyzed with GraphPad 9.4.1 Prism (GraphPad Software, San Diego, CA) and IBM SPSS 27 programs (IBM, Chicago, IL). The human data were analyzed using SAS 9.4 programs (SAS Institute Inc., Cary, NC) by a statistician who had no role in the clinical data collection or biospecimen processing. All hypothesis testing was 2-tailed and statistical significance was set at p≤0.05.

## 3. Results

### 3.1 Postoperative delirium-like behavior (Attention task) post-surgery in aged mice

Mice that underwent tibial fracture surgery had reduced attention performance in the 5-CSRTT when compared to naïve mice. There was a significant effect of the test day on the attention performance of the 5-CSRTT: [RM ANOVA, F(6,108) = 10.005, p < 0.001]. Further, there was a significant interaction between test day and surgery condition [RM ANOVA, F(6,108) = 3.800, p = 0.002], indicating that the reduced attention performance differed by both test day and whether the subject underwent surgery or naïve non-surgical controls. Examination of Bonferroni-corrected, pair-wise comparisons for each test day found that mice that underwent tibial fracture surgery had reduced attention performance in the 5-CSRTT on post-operative day 1 after surgery (p = 0.018, Mean Difference (MD)= -21.198, 95% CI [-38.391, -4.005]), day 4 (p = 0.046, MD= -11.265, 95% CI [-22.303, -0.226]) and on day 5 of testing (p=0.050, MD= -10.382, 95% CI [-20.755, -0.010]) (**Figure 1.A, Figure 1 sup.A**).

**Figure 1:**
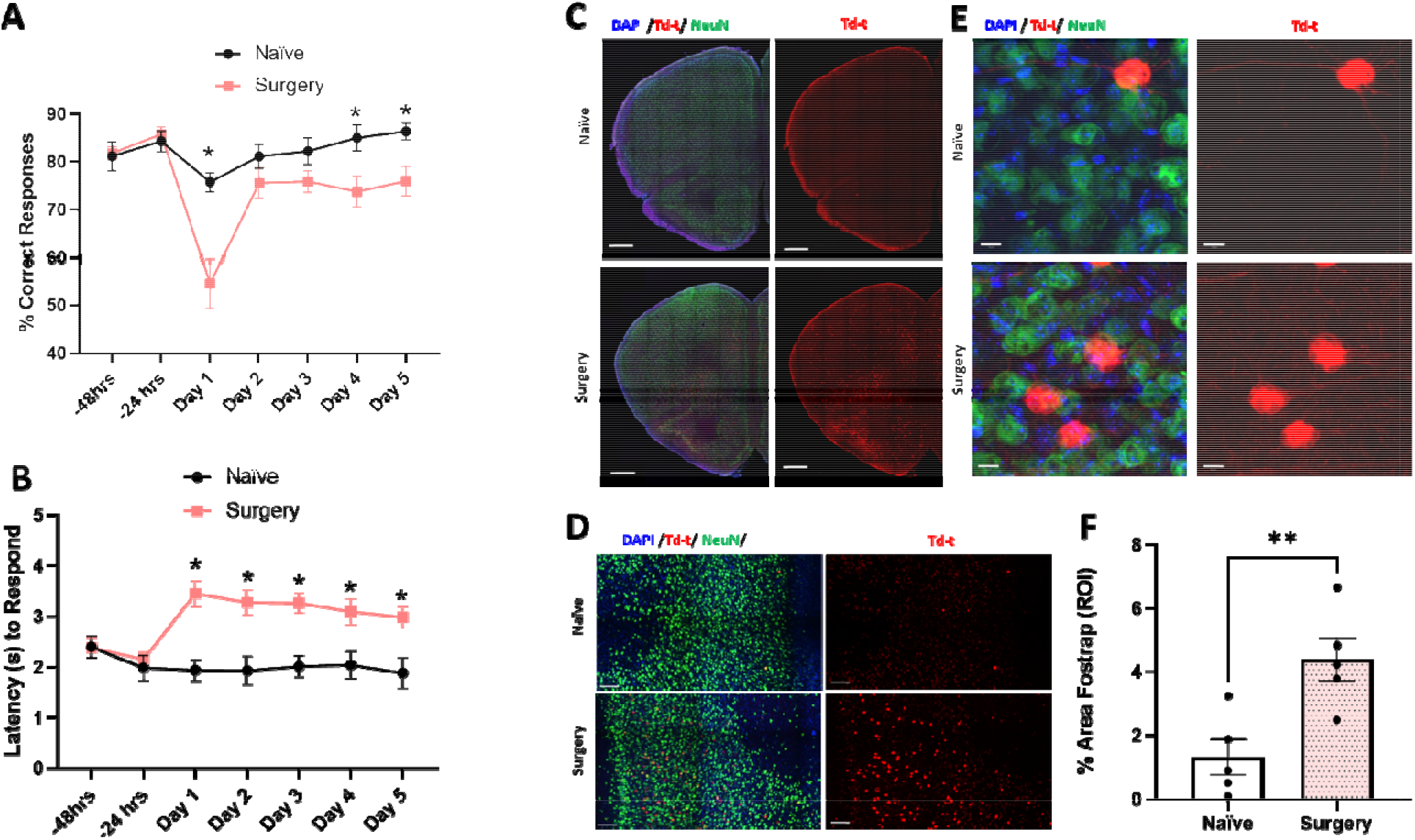
Delirium-like behavior and neuronal changes associated with surgery. Aged mice were trained in the 5-CSRTT, an attention task. A) Surgery revealed an acute deficit in attention processes at 24 hours and at 4 and 5 days post-tibial fracture. B) The latency to respond to the light stimulus was also examined and increased after surgery. C, D, E) Representative pictures of FosTRAP activation in FosTrap2 mice after surgery and naïve control mice showed in half brain, the PLC and with higher magnification of the PLC. F) Quantification data from panel D revealed that surgery increases FosTRAP activation in the PLC when compared to naïve control. Data are expressed as mean ± SEM. Analysis were performed as followed: RMANOVA, Bonferroni’s post hoc analysis and Unpaired Student’s T-test, * p < 0.05, ** p <0.01, ****p < 0.0001. Scale bars: 500 μm for C), 100 μm for D) and 10 μm for E).

In addition, as another index of performance, we also examined the latency to respond to the light stimulus in the 5-CSRTT. Here, the mice that underwent surgery took longer to respond across all post-treatment days in the task than the naïve non-surgical controls (**Figure 1.B, Figure sup 1.B**). There was a significant effect of the time (test day) on the latency to respond in the 5-CSRTT [RM ANOVA, F(6,108) = 2.281, p = 0.041]. Further, there was a significant interaction between test day and surgery condition [RM ANOVA, F(6,108)= 4.44, p= <0.001], indicating increased postoperative response latency in the surgery group but unchanged response latency in the naïve group. Bonferroni-corrected pair-wise comparisons between naïve control and surgery mice for each test day showed significant differences on post-operative test day 1 (p= 0.002, MD = 1.516, 95% CI [0.635, 2.398]), day 2 (p = 0.004, MD = 1.349, 95% CI [0.476, 2.222]), day 3 (p = 0.001, MD= 1.254, 95% CI [0.555, 1.953]), day 4 (p = 0.026, MD = 1.052, 95% CI [0.138, 1.967]) and day 5 of testing (p = 0.008, MD = 1.106, 95% CI [0.326, 1.885). Importantly, no group differences were detected at the two baseline days. Hence the mice that underwent tibial surgery were impaired on the several days post-surgery according to the percent correct responses and their latencies to respond to the light stimulus were greatly prolonged compared to the naive controls.

### 3.2 Neuronal activity changes after surgery

To determine which brain areas were most affected by the surgery, we used FosTRAP mice. Notably, an examination of FosTRAP expression revealed that activation of the prefrontal area and particularly the prelimbic cortex (PLC), a key region involved in attentional processing [23, 24], was strongly activated in mice subjected to the tibial surgery (**Figure 1.C**). Higher magnification showed that Td-tomato+ staining co-localized with NeuN+ staining, confirming that surgery activated Fos expression in neurons of the prelimbic area (**Figure 1.D-E**). Indeed, quantification of Td-tomato expression (*i*.*e*., Fos+ cells) in the medial prelimbic area supported a significant increase in in the surgical mice: Naïve: Mean: 1.317 (SEM: 0.5665); Surgery: Mean: 4.395 (SEM: 0.6852); Student’s T-test; (P = 0.0085; MD = 3.078, 95% CI [1.028, 5.128]) (**Figure 1.F**). Interestingly, no significant treatment effect was observed in the hippocampus (data not shown).

### 3.3 Astrocyte impairment and YKL-40 expression in the CNS after surgery in mice

We and others have previously described glial changes, particularly microglial activation after surgery [10, 11]. Here we focused on astrocytes, the most abundant cell type in the brain. Note, this cell type has not been extensively examined in this model. Expression of the astrocyte marker GFAP is known to be low in naïve conditions in the PLC [25] was also found to be the same in the present study (data not shown), immunohistochemical evaluation was not possible in this brain region. Due to this limitation, we assessed changes in GFAP and YKL-40 in the hippocampal dentate gyrus (**Figure 2.A)**.

**Figure 2:**
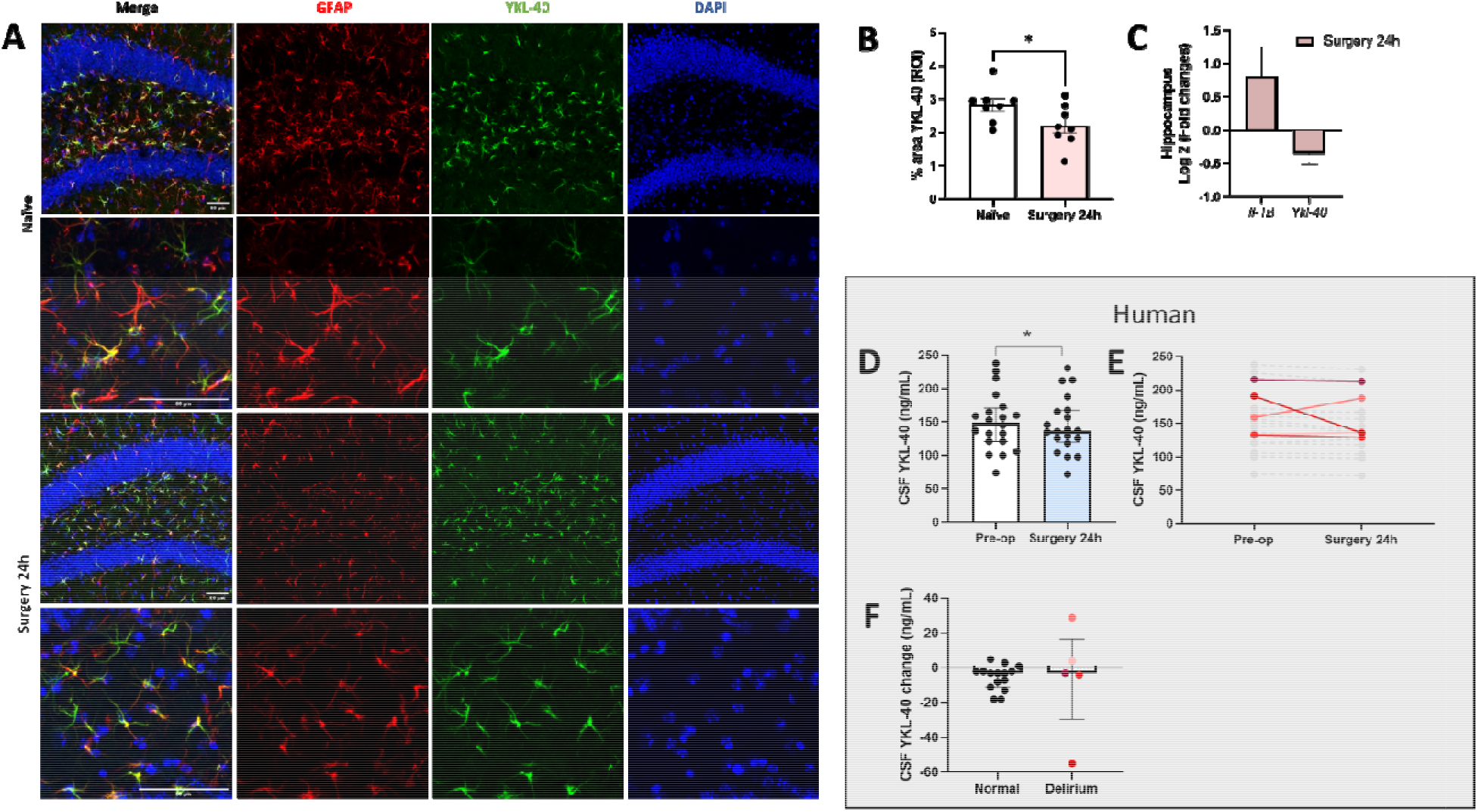
CNS YKL-40 changes following surgery in mice and human. Mouse brains were collected from aged mice 24 hours following tibial fracture or in naïve unoperated control mice. A) Representative immunohistochemical pictures of YKL-40 protein expression in the dentate gyrus of the hippocampus. B) The percent area of YKL-40 immunostaining was lower in the tibial surgery mice than naïve controls, C) RT-PCR of *Ykl-40* mRNA expression expressed as log2 of fold changes in hippocampus found *Ykl-40* levels were reduced following tibial fracture. Interestingly, RT-PCR *Il1-B* log 2 of fold change in hippocampus analysis showed increased *Il1-B* levels following tibial fracture. D) YKL-40 levels in CSF samples from human surgical patients examined prior to and 24 hours after surgery revealed a significant decrease in YKL-40 in the CSF following surgery in these patients. E) Individual patient’s pre-operative and post-operative levels are represented in grey (normal) or colored (patients with delirium), F) Patients were divided in two groups: normal and patients with delirium. CSF YKL-40 change levels (24 hours versus pre-operative) between normal and patients with delirium. Mice data are expressed as mean ± SEM and human data as mean ± SD except for CSF YKL-40 Median ± [interquartile range]. Analysis were performed as followed: Unpaired Student’s T-test, Wilcoxon Rank sum test when appropriate, * p < 0.05. Representative scale for A) is 50μm. Note: Individual dot colors represent the same patient with delirium across panels E and F.

At 24 hours after surgery, YKL-40 protein immuno-reactivity expression in the dentate gyrus was significantly decreased in the mice that had surgery relative to controls: Naïve: Mean: 2.826 (SEM: 0.1888); Surgery: Mean: 2.198 (SEM: 0.2202); Student’s T-test; (P = 0.0481; MD = - 0.6281, 95% CI [-1.25, -0.006]) (**Figure 2.B**). *Ykl-40* mRNA expression was also examined and it was found to be decreased at this time point by -0.355 log 2 (fold changes) (**Figure 2.C)**.

To corroborate the loss of YKL-40 expression and due to its regulation by the macrophage-released pro-inflammatory mediators Interleukin-1β (IL-1β) [26] we also measured expression of *Il-1-B* mRNA to confirm neuro-inflammation, which increased by 0.806 log2 (fold changes) as previously described in this model (**Figure 2.C**).

### 3.4 YKL-40 changes in human CSF after surgery

To determine whether our results with YKL-40 in mice could be replicated in humans, we assayed the levels of this protein in the CSF of human patients undergoing surgery. Overall YKL-40 levels in the human CSF were reduced 24 hours after surgery relative to the levels in the same patients before surgery (median Pre-op: 149.2 ng/ml [Q1:121.1, Q3:169.1]; 24 hours post-op: 136.7 ng/ml [Q1:121.9, Q3:166.8] Wilcoxon Rank Sum Test; p = 0.030) (**Figure 2.D**), and when considering individual subject trajectories pre- and post-surgery (**Figure 2.E**). However, there was no difference in the post-surgical change between patients with or without delirium, (median [Q1, Q3] with -3.2 [-4.0, 3.6] versus without -3.3 [-11.3, -1.7] delirium Wilcoxon Rank Sum Test; p = 0.601) (**Figure 2.F**).

### 3.5 Increased plasma YKL-40 after surgery both in mice and humans

We then measured YKL-40 in plasma samples that were drawn at the same time as the CSF samples, from the same human subjects. In human plasma, YKL-40 levels increased from before to 24 hours following surgery, (Pre-op: Mean: 72.3 ng/ml (SD: 76.6); 24 hours post-op Mean: 279.7 ng/ml (SD: 203.3); Student’s T-test; p = 0.001) (**Figure 3.A,B**). In addition, we observed a potential trend towards greater increase in YKL-40 from pre-op to 24-hours post-op among patients with delirium compared to patients without delirium. The average YKL-40 increase (SD) for those without delirium was 156.7 ng/ml (SD: 164.7) versus 336.2 ng/ml (SD: 277.6) for those with delirium, Student’s T-test; p = 0.114 (**Figure 3.C**). Similarly, mouse YKL-40 plasma levels at 24 hours were higher following tibial fracture versus sham treatment: Naïve: Mean: 26.55 ng/ml (SEM: 2.265); Surgery: Mean: 50.36 ng/ml (SEM: 4.153); Student’s T-test; (P = 0.0002; MD = 23.81, 95% CI [14.18, 33.45]) (**Figure 3.D)**. Together with YKL-40, we also observed elevated plasma IL-6: Naïve: Mean: 8.828 pg/ml (SEM: 4.358); Surgery: Mean: 55.10 pg/ml (SEM: 4.879); Student’s T-test; (P < 0.0001; MD = 46.28, 95% CI [32.02, 60.53]) (**Figure 3.E**), which is consistent with prior work using this mouse model and in a study of human plasma from patients with delirium [27, 28].

**Figure 3:**
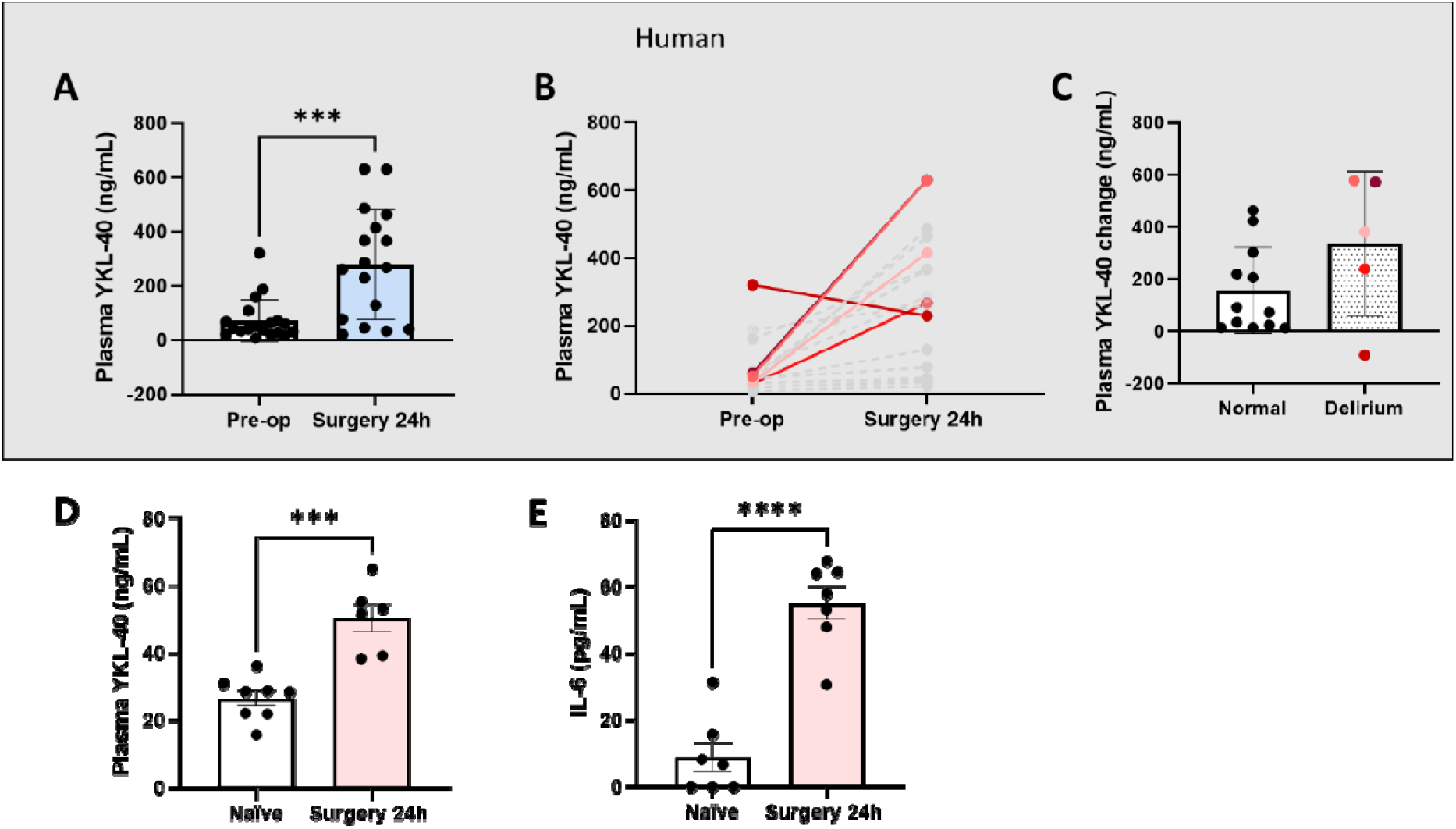
Plasma YKL-40 changes following surgery in humans and mice. A) Human plasma concentrations of YKL-40 revealed increased levels following surgery when compared to pre-operative levels within the same patients; B) Individual patients pre-operative and post-operative levels are represented in grey (normal) or colored (patients with delirium). C) A trends toward increase was observed in plasma YKL-40 change levels (24 hours versus pre-operative) between normal and patients with delirium. D) Mice plasmas were collected from aged mice 24 hours following tibial fracture or in naïve control mice. Similar increase in mice plasma YKL-40 was observed 24 hours following surgery. E) Plasma IL-6, a marker of inflammation revealed that aged mice had increased inflammation 24 hours after surgery. Human data are expressed as mean ± SD and mice data as mean ± SEM. Analysis were performed as followed: Paired and Unpaired Student’s T-test when appropriate, ***p < 0.001. Note: Individual dot colors represent the same patient with delirium across panels B and C.

### 3.6 Changes in NfL and BBB opening after surgery in mice

Using the same murine plasma samples, we performed SIMOA analyses of NfL. Given the peak of inattention and established neuroinflammation on post-operative day 1, we evaluated NfL at this time Plasma levels of NfL were significantly increased 24 hours after tibial fracture: Naïve: Mean: 153.3 pg/ml (SEM: 15.94); Surgery: Mean: 755.1 pg/ml (SEM: 57.36); Student’s T-test; (P < 0.0001; MD = 601.8, 95% CI [472, 731.5]) (**Figure 4.A**). Albeit we did not investigate the origin of NfL in the current study, we observed that the BBB was significantly impaired in the aged mice after surgery, suggesting a possible way for brain-derived NfL to enter the bloodstream. In fact, loss of mRNA key tight junction markers was evident as early as 9 hours after surgery and was still impaired 24 hours post tibial fracture surgery (*occludin, Cd31, Zo1, claudin5* and *Aq-4*) by RT-PCR (**Figure 4.B**).

**Figure 4:**
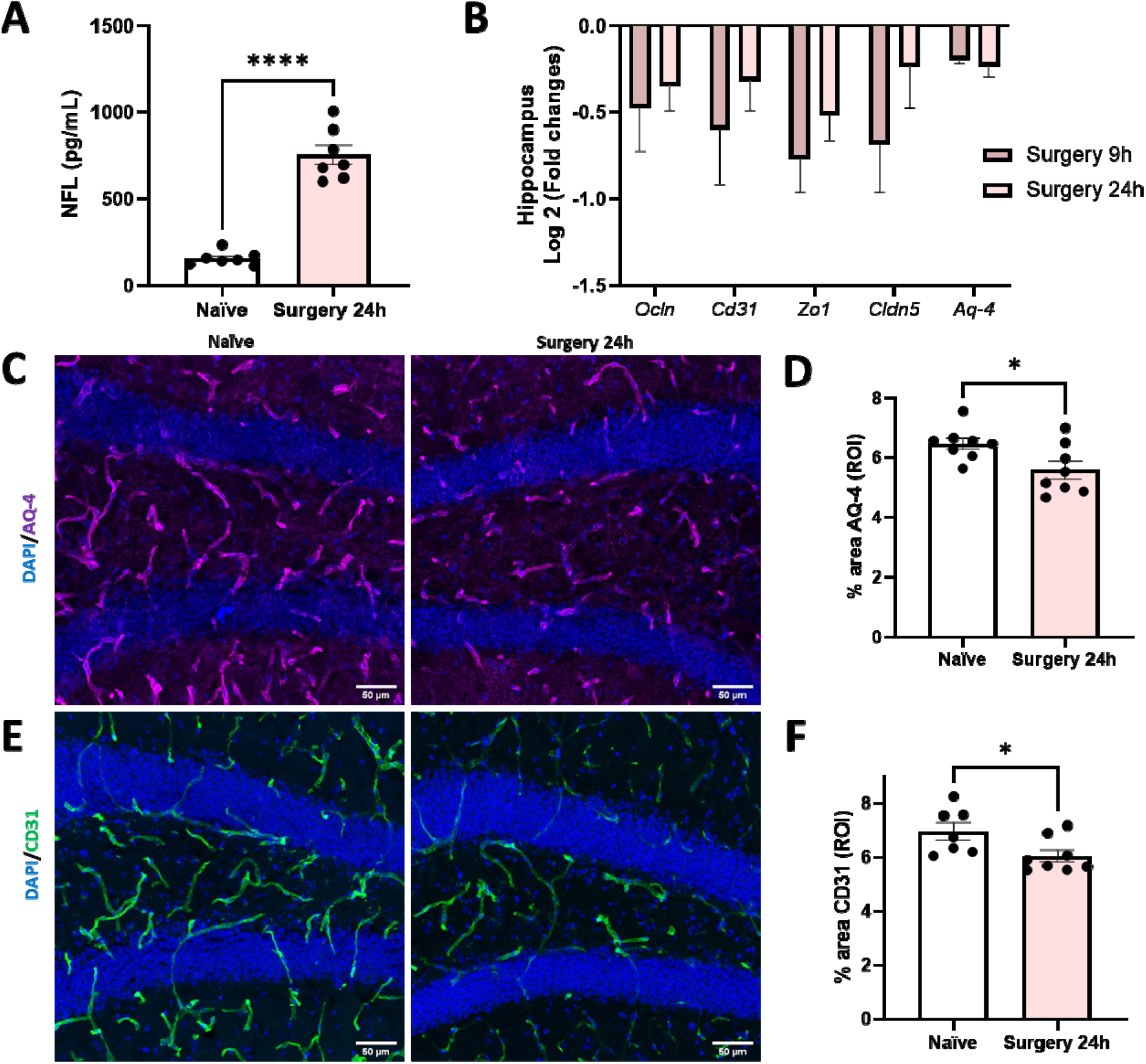
BBB impairments following surgery. Mice brains were collected from aged mice 24 hours following tibial fracture or in naïve control mice. A) Using Quanterix Simoa, Plasma NfL, a marker of neuronal damage revealed that aged mice had increased levels 24 hours after surgery B) RT-PCR log 2 of fold changes of *Occludin, Cd31, Zo1, Claudin5* and *Aq-4* revealed BBB impairment at 9 and 24 hours following tibial fracture. C) Representative pictures of the astrocytic endfeet AQ-4 and E) the endothelial cell marker CD31 revealed decreased percentage area covered by D) AQ-4 and F) CD31 staining in the dentate gyrus following orthopedic surgery. Data expressed as mean ± SEM. * p < 0.05, Analysis were performed as follows Unpaired Student’s T-test. Representative scale for C, E) is 50μm.

In addition, we used immunohistochemistry to evaluate expression of the astrocytic endfoot and BBB integrity marker AQ-4 and the endothelial cell marker CD31. AQ-4 protein immuno-reactivity expression in the dentate gyrus ROI was decreased 24 hours after surgery: Naïve: Mean: 6.473 (SEM: 0.197); Surgery: Mean: 5.587 (SEM: 0.2955); Student’s T-test; (P = 0.0258; MD = -0.8854, 95% CI [-1.647, -0.1237]) (**Figure 4.D)**. Similarly, CD31 protein immuno-reactivity expression was reduced at the 24 hour time point: Naïve: Mean: 6.958 (SEM: 0.3183); Surgery: Mean: 6.05 (SEM: 0.2283); Student’s T-test (P = 0.0346; MD = -0.9074, 95% CI [- 1.738, -0.07679]) (**Figure 4.F**).

## 4. Discussion

This project aimed at comparing neuroimmune endpoints relevant to delirium pathophysiology both in mice and humans after surgery. We described a conserved YKL-40 response to anesthesia and surgery in mice and humans, with an increase in plasma levels but a decrease in CNS levels. Notably, evaluation of YKL-40 expression in the mouse brain revealed a significant decrease of YKL-40 in hippocampal astrocytes after surgery. This raises the possibility that a similar decrease in CNS YKL-40 expression may occur following surgery in humans, which would fit with the decrease in YKL-40 levels in the human CSF we observed following surgery. In mice, we also found a significant increase in plasma NfL after surgery, which was associated with BBB breakdown. Overall, these data validate the usefulness of animal models to clarify neuroimmune mechanisms in postoperative delirium. Further, these results advance our understanding of the dysfunctional cellular and molecular processes that may contribute to postoperative delirium pathogenesis in the aging brain.

This study leveraged complementary multi-species assays to advance our understanding of postoperative delirium. In fact, by using a subset of the well-phenotype INTUIT cohort with parallel plasma and CSF analyses together with a mouse model of orthopedic surgery, we have described remarkable similarities in the neuroimmune response to anesthesia and surgery. We have particularly focused on YKL-40 due to its growing role in AD neurodegeneration as well as postoperative delirium. Recently, Vasunilashorn *et al*. [16] described plasma YKL-40 as a key delirium-associated protein both as a risk marker of delirium (*i*.*e*., preoperative elevation) and as a disease marker increased more in delirious patients on postoperative day 2 than non-delirious patients. In their proteome-wide analysis of plasma in 18 older orthopedics surgery patients with postoperative delirium and 18 matched controls, Vasunilashorn *et al*. identified YKL-40 as the only protein of > 1,300 evaluated whose preoperative elevation predicted delirium and whose postoperative elevation trend followed delirium onset and, possibly, resolution [16]. These findings suggesting that YKL-40 may be a useful plasma biomarker for delirium are not surprising as YKL-40 has been associated with ICU mortality in sepsis [29] prion disease, [13, 30] Alzheimer’s disease, [13, 29] and other dementias (vascular, Lewy body, fronto-temporal), ; however, the plasma YKL-40 levels cannot definitively link central YKL-40 to delirium. We found plasma YKL-40 changes are consistent with prior reports in our clinical cohort, albeit we were underpowered to detect changes in delirious subjects. Based on the level of variability in the 24h change in plasma YKL-40 observed in the human samples (212 ng/ml), a future human study with a 12% delirium incidence would require **297** subjects (36 with and 261 without delirium) to detect a Cohen’s effect size of 0.50. Based on the 24 hours plasma YKL-40 increase of 156.7 ng/ml in our human subjects without delirium, such a study would be able to detect a 1.68 times higher YKL-40 increase (263 ng/ml) among those developing delirium

Interestingly, we found a similar response in aged mice after surgery, suggesting that YKL-40 increase in the periphery is likely a conserved response to tissue injury possibly mediated by peripheral immunocytes [31]. However, for both mice and humans the YKL-40 response in the CSF was surprising and the opposite of what we found in plasma. In fact, CSF YKL-40 levels were lower 24 hours after surgery, with no detectable difference between the subjects that had delirium. In [15] no changes were observed in CSF YKL-40 postoperative, possibly explaining a different etiology than other neurodegenerative conditions with robustly elevated YKL-40 in the CSF [13, 14, 29, 30, 32-37]. In the CNS astrocytes are the primary source of YKL-40, which provides an attractive target for AD [38].

Here we evaluated astrocytes pathology in the mouse hippocampus and confirmed the reduction of YKL-40 expression both at mRNA and protein levels 24 hours after orthopedic surgery. At this time point GFAP+ astrocytes were visibly dystrophic and YKL-40 expression mostly was confined within these cells. It is possible that YKL-40 requires more time to become upregulated in the CNS, given that delirium is characterized by a rapid and acute onset. Astrocyte dysfunction in this model was previously described [39], suggesting a role not only for neuroinflammation but also for neurometabolic dysfunction in delirium. In fact, levels of *Il-1-B* mRNA expression were increased despite *Ykl-40* mRNA expression being reduced at 24 hours, possibly contributing to prolonged astrocyte pathology in the vulnerable brain [40, 41]. Overall, the murine evidence suggests that the postoperative human decrease in CSF YKL-40 levels truly reflects central processes.

Astrocytes are also important regulators of the BBB [42]. In this study, we confirmed the loss of astrocytic-end-feet processes and endothelial markers at 24 hours after surgery in mice, suggesting BBB opening is a major culprit for ensuing delirium-like behavior. Here we confirmed the validity of another plasma biomarker found upregulated in delirious subjects: NfL [43, 44]. The abundance of NfL in plasma together with early signs of BBB opening may indicate a central source for this neuronal injury marker. The ability to interrogate CNS damage via systemic biofluids is highly tractable in the clinic, however, peripheral nerve damage itself could contribute to the cellular source of NfL in plasma. Further characterization of this cellular response is needed not only in delirium but also AD and other ADRDs.

Importantly, these data add to the frameworks that delirium biomarkers are dynamic. These dynamic biomarkers may not represent global cognitive changes as they manifest in this syndrome. For example, BBB changes are limited to select areas of vulnerability, suggesting that only particular regions of interests are susceptible to anesthesia and surgery. This explains why molecules, including peripheral YKL-40, do not passively diffuse across the permeable BBB. While plasma biomarkers are undoubtedly more accessible and feasible in human clinical medicine than CSF biomarkers, they must be interpreted cautiously. This need for caution is apparent in our findings that plasma YKL-40 increases while CSF and murine data suggest a central YKL-40 decrease. One means to guide and enrich plasma biomarker interpretation is to evaluate these markers in alternative models, such as with parallel human CSF analysis and/or the trans-species model with murine histology and RT-PCR analysis presented here. Overall, the differential change observed in plasma and CSF YKL-40 suggests cautious use of plasma biomarkers that may not specifically reflect brain changes. More broadly, this study encourages evaluation of possible alternative sources of plasma biomarkers and provides rational about the importance of evaluating BBB integrity. Indeed, these results suggest BBB breakdown and crosstalk between the periphery (systemic sources) and the CNS means by which cytokines may cross the BBB and facilitate inflammation inside the brain.

Further, we demonstrated ways to study delirium mechanisms using parallel human and rodent models. For example, the attention deficit characterized in the 5-CSRTT in mice is homologous to the inattention delirium sub-feature observed in human patients and facilitates translational study between mice and humans [10]. The present study demonstrates this; that animals subjected to tibial fracture have decreased performance in 5-CSRTT within 24 hours after surgery compared to intact controls. Moreover, despite the surgerized mice exhibiting a recovery exemplified by increase success rates over subsequent test days, these animals remain below the performance levels of intact animals at the same time points. This small reduction in success rates observed during the recovery period is further compounded by sustained increases in the delayed latency to respond in the task. Further, we report increased post-tibial fracture surgery plasma NfL levels in mice, which parallels similar reports in delirious older patients after surgery [43, 44]. In humans, there are also limitations to studying neuronal activation at a cellular level; however, in this study using FosTRAP mice, we showed post-tibial fracture PLC neuronal activation in mice. This result links the behavior observed in the human delirium phenotype (decreased attention) with a potential neuronal mechanism (PLC neuronal activation) in mice that can only practically and ethically be performed in animal models. Collectively, this cross-species evaluation allowed us to use histology to fill in the gaps necessitated by human biomarker studies while grounding our pre-clinical models with parallel data samples from human patients.

## Limitations

There are a few limitations to this study. The breeding time required for FosTRAP mice and the complexity of Tamoxifen-induced recombination (e.g, successive daily injections required in older mice) made the use of older mice for the FosTRAP experiments impossible for this specific study. Attentional and delirium evaluations in both mice and humans focused on the 24-hour time point where inattention peaked in the mouse 5-CSRTT and where all human subjects were still hospitalized and, thus, available for 3D-CAM assessment. Future studies should characterize the long-term cognitive sequelae of surgery and/or delirium. We used a limited number of human patients with versus without delirium to confirm our findings in mice. The available human samples came from patients who underwent a variety of non-cardiac, non-neurological surgery, only some of which were orthopedic. In addition, human patients were not chosen randomly as they were selected for *APOE4* carrier status in conjunction with a separate project. Therefore, there is an overrepresentation of *APOE4* carriers in this cohort. Finally, the human subjects in this study underwent various types of surgery while the rodent model only replicated orthopedic injury / surgery. That said, all surgeries have inherent musculocutaneous injury with incision, and the conserved responses observed despite different surgery types imply that sufficient homology is present between species. Thus, cross-species postoperative delirium models likely do not require identical rodent and human surgeries, just as clinical delirium research often includes patients undergoing many different surgery types.

## Conclusion

In this study, we confirmed that plasma biomarkers associated with human delirium are also elevated in mice postoperatively. We built on those human findings with a more in-depth, histological analysis of transgenic FosTRAP mice and with mRNA expression assessments. Human retrospective studies have shown that patients who develop postoperative delirium or sustain other neurological injuries (*e*.*g*., traumatic brain injury) tend to have higher levels of plasma biomarkers associated with astrocyte function, general inflammation, and neuronal injury. In this study, we showed that levels of the plasma inflammatory marker IL-6 increased at 24 hours post-tibial fracture surgery in mice. Although we did not measure IL-6 in humans here, given that inflammation occurs after any trauma including surgery, and that massive systemic inflammation in sepsis often co-occurs with delirium, previous studies revealed that the inflammatory marker IL-6 is significantly higher in the plasma of postoperative patients who develop delirium compared with those who do not [45, 46]. Overall, this cross-species study using complementary mouse and human preparations yield novel insights that neither a rodent nor human study could produce independently. We demonstrated that there is a conserved YKL-40 response, increases peripherally and decreases centrally, among aged mice following orthopedic surgery and older human patients. These findings urge cautious interpretation of plasma delirium biomarkers considering evidence of BBB compromise and unanticipated peripheral biomarker sources.

## Acknowledgements

We would like to acknowledge the assistance of the Duke Molecular Physiology Institute Molecular Genomics Core for the generation of the NfL data (RRID:SCR_017860); the Duke Behavioral Core (Christopher Means); the Duke Microscopy core (Lisa Cameron and Benjamin Carlson), and Fan Wang (MIT) for her gift of the Fos-TRAP mice.

NT acknowledges support from National Institutes of Health grants R01-AG057525 and P01AT009968-A1, Alzheimer’s Association (2019-AARG-643070) and the Anesthesiology Department. HZ is a Wallenberg Scholar supported by grants from the Swedish Research Council (#2018-02532), the European Union’s Horizon Europe research and innovation programme under grant agreement No 101053962, Swedish State Support for Clinical Research (#ALFGBG-71320), the Alzheimer Drug Discovery Foundation (ADDF), USA (#201809-2016862), the AD Strategic Fund and the Alzheimer’s Association (#ADSF-21-831376-C, #ADSF-21-831381-C, and #ADSF-21-831377-C), the Bluefield Project, the Olav Thon Foundation, the Erling-Persson Family Foundation, Stiftelsen för Gamla Tjänarinnor, Hjärnfonden, Sweden (#FO2022-0270), the European Union’s Horizon 2020 research and innovation programme under the Marie Skłodowska-Curie grant agreement No 860197 (MIRIADE), the European Union Joint Programme – Neurodegenerative Disease Research (JPND2021-00694), and the UK Dementia Research Institute at UCL (UKDRI-1003). LA acknowledges support from NIH T32GM008600.

## Author’s contributions

JD, LA, AC, YW, SG performed experiments and analyzed data, NF, RMR, WW conducted and analyzed the behavioral study, MCW overviewed the statistical analyses, MD, HZ, TY contributed key reagents, MB supervised the human study, NT, JD and LA conceived the idea. JD, LA, MB, NT wrote the manuscript with input from all authors. All authors read and approved the manuscript.

## Figures

**Figure 1 Supplementary:**
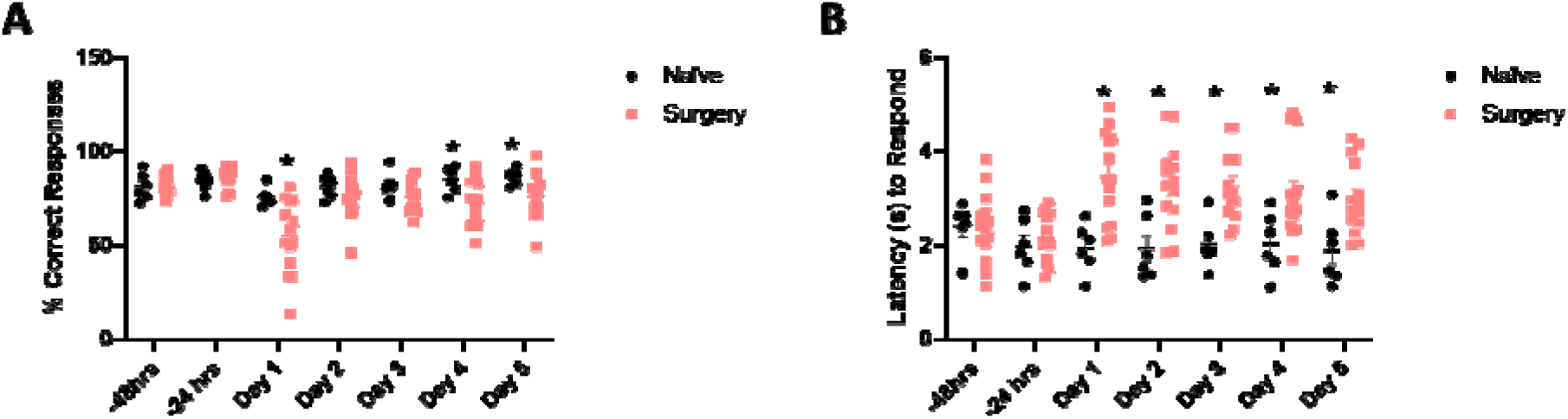
Delirium-like behavior, individual mouse data. Aged mice were trained in the 5-CSRTT, an attention task. A) Surgery revealed an acute deficit in attention processes at 24 hours and at 4 and 5 days post-tibial fracture, B) The latency to respond to the light stimulus was also examined and increased after surgery.

